# On the expression of reproductive plasticity in *Drosophila melanogaster* females in spatial and socially varying environments!

**DOI:** 10.1101/2023.12.19.571982

**Authors:** Oghenerho Nwajei, Sanduni Talagala, Laura Hampel, Bhavya Punj, Nuek Yin Li, Tristan A.F. Long

## Abstract

Individuals often adjust their behaviour based on their perception and experiences with the social and/or physical environment. In this study, we examined the extent of reproductive plasticity expressed in mating rates, mating latencies, mating durations, and offspring production in female fruit flies, *Drosophila melanogaster*, that encountered different numbers of males in different sized chambers. We found that mating latency length decreased with more courting males and smaller environments and that matings durations were longer in larger chambers and in the presence of two males. These results illustrate the sensitivity of these behavioural phenotypes to changes in local environmental conditions.

## Discussion

Fruit flies, *Drosophila melanogaster*, are frequently used in studies designed to explore how genetic and/or social factors affect the expression of pre- and post-copulatory sexual selection and sexual conflict. The courtship and mate assessment in this species is complex and involved the exchange of visual, auditory, chemical and tactile cues (Welbergen et al. 1992, Hall 1994, Greenspan & Ferveur 2000). By examining the frequency of matings, how long it takes for flies to start mating (“mating latency”), and the duration of copulation (“mating duration”), in different experimental scenarios, the plastic responses of the sexes can be used to infer how males and females perceive their mates and their reproductive environment (Friberg 2006, Bretman et al. 2009, Taylor et al. 2013). In our study we set out to explore how manipulating the number of males present in a mating arena (1 or 2) and the relative size (large or small) of said arena, influenced reproductive behaviour and subsequent offspring production.

While mating rate did exhibit a marginally significant interaction between the number of males and the chamber size treatments (Table 1a), post-hoc comparative tests did not reveal any significant differences between pairs of treatment combinations. Mating was observed in 96.2% of vials in which there were 2 males, and 88.8% of vials with one male, a difference that was not statistically significant at the α=0.05 level, and of negligible effect size (Cliff’s delta & [95%CI] = 0.066 [0.136, 0.004]). While the overall rate of matings did not differ meaningfully between different social or physical environments, the temporal patterns of mating did. Female flies housed with 2 males mated faster (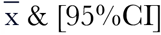: 1173s [976, 1411]) than did those housed with a single male (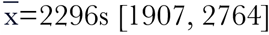; Cliff’s delta= 0.412 [0.273, 0.534]; Figure 1a & b, Table 1b), a result consistent with previous work that indicates that male courtship of females is higher when there is an increased risk or intensity of sperm competition (Bretman et al. 2009). Furthermore, mating resulted more quickly in the small chambers 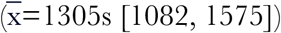 than in the large chambers (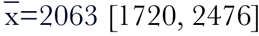; Cliff’s delta=0.406 [0.265, 0.530]), a result likely due to the increased ease of males locating, pursuing and courting females in the smaller environment. Previous studies have found that males engage in more intensive courtship for females in smaller space spaces (Ewing and Ewing 1984), likely because it is more difficult for females to flee from unwanted male courtship in a smaller arena (Partridge et al. 1987).

**Table 1.** Analysis of deviance tables examining the effects of size of chamber size, numbers of males present in the chamber and their interaction on (a) the likelihood of mating during the observation session (b) the latency to mate, (c) the duration of subsequent copulations, and (d) the number of offspring produced in the next 24h in *D. melanogaster*.

| Factors | a) Mating Rate |  |  | b) Latency |  |  | c) Duration |  |  | d) Offspring |  |  |
| --- | --- | --- | --- | --- | --- | --- | --- | --- | --- | --- | --- | --- |
| | LR $\chi^2$ | df | p | LR $\chi^2$ | df | p | LR $\chi^2$ | df | p | LR $\chi^2$ | df | p |
| Chamber Size | 0.002 | 1 | 0.966 | 11.747 | 1 | 0.001 | 4.020 | 1 | 0.044 | 8.564 | 1 | 0.003 |
| Number of Males | 3.512 | 1 | 0.061 | 25.330 | 1 | 4.8x10<sup>-7</sup> | 5.521 | 1 | 0.018 | 3.007 | 1 | 0.083 |
| Size x Number | 4.144 | 1 | 0.042 | 2.043 | 1 | 0.152 | 2.183 | 1 | 0.140 | 0.060 | 1 | 0.806 |

**Figure 1.**
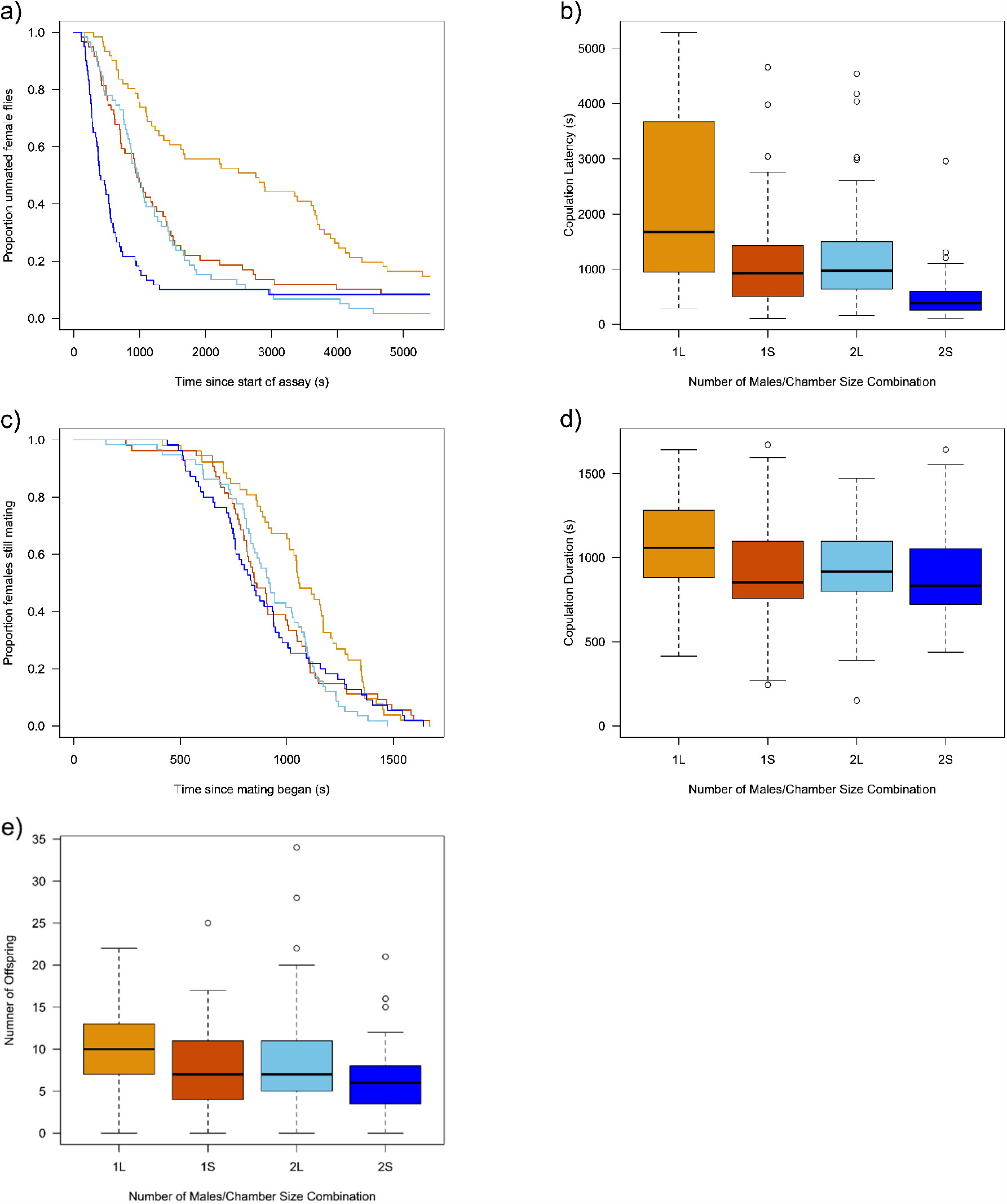
Reproductive behavioural plasticity and offspring production of female flies housed with either 1 or 2 males in different sized chambers (Small “S” or Large “S”). The phenology and distribution of mating latencies are depicted in Fig 1a & 1b; mating durations are plotted Fig 1c & 1d, and offspring production is depicted in Fig 1e For boxplots, the boxes enclose the middle 50% of data (the inter-quartile range, IQR), with the thick horizontal line representing the location of median. Data points > ±1.5*IQR are designated as outliers. Whiskers extend to largest/smallest values that are not outliers, indicated as closed circles. Colours used to differentiate different treatments in survivorship plots correspond to those used in the boxplots.

Mating duration also exhibited plasticity in its expression (Figures 1c & 1d). When two males were present in a vial, the length of copulation was shorter 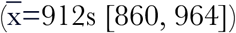 than when there was only the single mating male (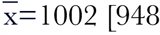, 1055; Cliff’s delta=0.169 [0.015, 0.315]; Table 1c). This phenomenon has also been seen in a previous study (Bretman et al. 2009) where it was hypothesized that is was due to interference from the unmated male, or a case of strategic adjustment by the mating male in anticipation of encountering sperm competition in the future. Alternatively, as females do play an active role in shaping the length of copulation (Bretman et al. 2013, Edward et al. 2014) they may be more motivated to end mating sooner in the presence of another potential mate. Interestingly, the acoustic signal of male courtship songs triggers an escape response in females (Arez et al. 2021), so if the second male courts the mating female (something that occurs frequently -*see* Dukas 2020) this might conceivably lead to in an increased chance of earlier termination. Matings that occurred in smaller chambers were marginally shorter 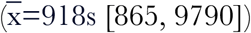 than those that happened in the larger arenas (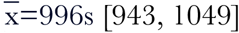; Cliff’s delta=0.202 [0.050, 0.345]), possibly as it was more difficult for females to dislodge males in the more confined area.

When examining offspring production, females in larger chambers produced more offspring on average 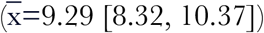 than those in small chambers (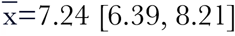; Cliff’s delta= 0.237 [0.090, 0.374]). This small difference might be attributable those females in larger spaces experiencing less disruption to their offspring production and provisioning. In *D. melanogaster*, mated females must feed consistently to acquire the nutrients required to produce many yolk-rich ova (Spieth 1974), and persistent harassment by males may interfere with their ability to optimally forage (Bretman & Fricke 2019). Higher levels of mating and/or male harassment causes females to be less fit (Fowler and Partridge 1987, Partridge and Fowler 1990, Long et al. 2009). While there was no statistically significant difference in the number of offspring associated with the number of males present in the vial, there was a trend toward more offspring produced in the single male vials 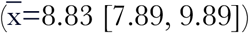 than in the vials with two males 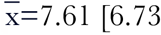, 8.60]; Cliff’s delta=0.182 [0.037, 0.319]) that is also consistent with the idea that greater potential exposure to males interferes with offspring production in females (Long et al. 2009).

Cumulatively, these results illustrate the considerable plasticity in the expression of some reproductive behavioural phenotypes in *D. melanogaster*, and underscores the need to consider both the social and the physical environment when designing experiments measuring their mating latencies and/or durations, as these are often used as proxies of individual attractiveness or are used to infer the costs and benefits of variation in the expression of reproductive behaviours (*see* Shackleton et al. 2005, Friberg 2006, Bretman et al. 2009, Edward et al 2014, Tennant et al. 2014)

## Methods

### Fly population and culture protocol

Our experiment used *Drosophila melanogaster* flies from the large (∽3500 adults/generation), outbred, wild-type population “Ives” (hereafter “IV”) that was founded from a sample of 200 male and 200 female flies that collected near South Amherst, MA (USA) in 1975, and have been cultured following a standardized protocol since 1980 (Rose 1984, Tennant et al. 2014). Flies in this population are maintained on non-overlapping 14-day generations, and develop in sets of 25×95 mm vials, (VWR: 75813-160) each containing ∽10mL of banana/agar/killed-yeast media as a light sprinkle of live yeast and are kept at 25°C, 60% relative humidity, and on a 12L:12D diurnal cycle. A standard density of 100 eggs/vial is established at the start of each culture cycle.

### Experimental Details

Adult male and female flies were collected as virgins (within 6 hours of their eclosion from pupae) from standard IV population vials. Females were placed into individual vials, and males were haphazardly placed into vials by themselves (“one male” treatment), or in same-sex pairs (“two male” treatment). Flies were left in these vials for ∽40h in order to fully mature prior to the start of the assay. The experiment began by transferring (without CO_2_ anesthesia) the female into the male vial. In half of the vials the plug (Droso-Plugs, Genesee Scientific: 59-200) was pushed down in the vial to create a mating area approximately 2cm in height (“small chamber” treatment), while in the other half, the plug was only pushed down in the vial sufficiently to create a 6cm high mating area (“large chamber” treatment). For each of the 4 treatment combinations we created ∽60 replicate vials. These vials were immediately mounted horizontally in a well-lit room and we carefully observed the flies for the next 90 minutes. If a copulation was observed during this period, the time that the event began and ended were both recorded to the closest second. From these observations the mating latency and mating duration were calculated. While it was impossible for the observers to be blind to the differences in the vial configurations or number of occupants, they were not told what the objective of the experiment was. Once the observation session was complete, the vials were left undisturbed for 24 hr before the flies were removed, and the vials we placed in an incubator for 13 more days. At that time the number of eclosed adult flies in each vial were counted.

### Statistical Analyses

All data analyses were conducted in the R statistical computing environment (version 4.0.3, R Core Team, 2020). We modelled the phenological changes in state for the mating latency and mating duration data using parametric regressions using the *survreg* function in the *survival* package (Therneau 2022, Therneau and Grambsch 2000). For the mating latency model, flies that did not mate by the end of the observation session were assigned the maximum value of 5400s, and right censored. In both models the the independent factors were the size of the chamber, the number of males present, and their interaction, and we selected the error distribution based on which yielded the lowest residual deviance (Crawley 2005). Consequently, we chose a Weibull error distribution for the mating latency model and a gaussian error distribution for the mating duration model. Both the frequencies of mating and the number of offspring were analyzed using General Linear Models (GLMs) where the independent factors were the size of the chamber, the number of males present, and their interaction. For the mating rate GLM, we used a binomial error distribution, while in the offspring GLM we employed a quasipoisson error as the data were over-dispersed (Crawley 2005). We tested the significance of independent factors in all models using a likelihood-ratio Chi-square test using the *Anova* function in the *car* package (Fox and Weisberg 2019). We estimated the marginal mean values (and 95%Cs) for different treatment using the *emmeans* function in the *emmeans* package (Lenth 2022). We also quantified the magnitude of the difference between groups of flies in different treatments with the Cliff’s delta effect size statistic using the *cliff*.*delta* function in the *effsize* package (Torchiano 2020)

### Reagents

*Drosophila melanogaster* from the “Ives”/ “IV” population

## Acknowledgements

The initial inspiration for this experiment arose from a lab discussion of a paper by Anastasio et al. (2023). Michael Steeleworthy of the Wilfrid Laurier University Library is thanked for their help with data archiving. This work was conducted at Wilfrid Laurier University, which exists on the traditional territory of the Neutral, Anishnawbe, and Haudenosaunee peoples.

## Author Contributions

ON, ST, & TAFL conceived and planned the experiment. TAFL & OG pushed flies. Data collected by ST, LH, BP, NYL & OG. Data analysis & interpretation by ON & TAFL. Manuscript draft by TAFL., with edits by ST.

## Ethics

This study did not require approval from an ethics committee.

## Supporting Data

All data analysed in this manuscript are archived in Borealis, the Canadian Dataverse Repository: https://doi.org/10.5683/SP3/U8HNPU

## References

Anastasio, O.E., Sinclair, C.S. and Pischedda, A., 2023. Cryptic male mate choice for highquality females reduces male postcopulatory success in future matings. Evolution, 77(6), pp.1396–1407. doi.org/10.1093/evolut/qpad064

Arez, E., Mezzera, C., Neto-Silva, R.M., Aranha, M.M., Dias, S., Moita, M.A. and Vasconcelos, M.L., 2021. Male courtship song drives escape responses that are suppressed for successful mating. Scientific Reports, 11(1), p.9227. doi.org/10.1038/s41598-021-88691-w

Bretman, A., & Fricke, C. (2019). Exposure to males, but not receipt of sex peptide, accelerates functional ageing in female fruit flies. Functional Ecology, 33(8), 1459–1468. doi.org/10.1111/1365-2435.13339

Bretman, A., Fricke, C. and Chapman, T., 2009. Plastic responses of male Drosophila melanogaster to the level of sperm competition increase male reproductive fitness. Proceedings of the Royal Society B: Biological Sciences, 276(1662), pp.1705–1711. doi.org/10.1098/rspb.2008.1878

Bretman, A., Westmancoat, J.D. and Chapman, T., 2013. Male control of mating duration following exposure to rivals in fruitflies. Journal of Insect Physiology, 59(8), pp.824–827. doi.org/10.1016/j.jinsphys.2013.05.011

Crawley, M.J., 2005. Statistics: an introduction using R. John Wiley & Sons. Chichester.

Dukas, R., 2020. Natural history of social and sexual behavior in fruit flies. Scientific reports, 10(1), p.21932. doi.org/10.1038/s41598-020-79075-7

Edward, D.A., Poissant, J., Wilson, A.J. and Chapman, T., 2014. Sexual conflict and interacting phenotypes: a quantitative genetic analysis of fecundity and copula duration in Drosophila melanogaster. Evolution, 68(6), pp.1651–1660. doi.org/10.1111/evo.12376

Ewing, L.S. and Ewing, A.W., 1984. Courtship in Drosophila melanogaster: behaviour of mixed-sex groups in large observation chambers. Behaviour, 90(1-3), pp.184–202.jstor.org/stable/4534364

Partridge, L., Green, A. and Fowler, K., 1987. Effects of egg-production and of exposure to males on female survival in Drosophila melanogaster. Journal of Insect Physiology, 33(10), pp.745–749. doi.org/10.1016/0022-1910(87)90060-6

Fox, J. and Weisberg, S., 2019) An R Companion to Applied Regression, Third Edition. Sage, Thousand Oaks CA: https://socialsciences.mcmaster.ca/jfox/Books/Companion/

Friberg, U., 2006. Male perception of female mating status: its effect on copulation duration, sperm defence and female fitness. Animal Behaviour, 72(6), pp.1259–1268. doi.org/10.1016/j.anbehav.2006.03.021

Greenspan, R.J. and Ferveur, J.F., 2000. Courtship in Drosophila. Annual Review of Genetics, 34(1), pp.205–232. doi.org/10.1146/annurev.genet.34.1.205

Hall, J.C., 1994. The mating of a fly. Science, 264(5166), pp.1702–1714. jstor.org/stable/2883918

Lenth, R.V., 2022. emmeans: Estimated Marginal Means, aka Least-Squares Means. R package version 1.7.2, https://CRAN.R-project.org/package=emmeans

Long, T.A.F., Pischedda, A., Stewart, A.D. and Rice, W.R., 2009. A cost of sexual attractiveness to high-fitness females. PLoS Biology, 7(12), p.e1000254. doi.org/10.1371/journal.pbio.1000254

Partridge, L., Ewing, A. and Chandler, A., 1987. Male size and mating success in Drosophila melanogaster: the roles of male and female behaviour. Animal Behaviour, 35(2), pp.555–562. doi.org/10.1016/S0003-3472(87)80281-6

Partridge, L. and Fowler, K., 1990. Non-mating costs of exposure to males in female Drosophila melanogaster. Journal of Insect Physiology, 36(6), pp.419–425. doi.org/10.1016/0022-1910(90)90059-O

R Core Team, 2020. A language and environment for statistical computing. Vienna: R Foundation for Statistical Computing.

Rose, M.R., 1984. Laboratory evolution of postponed senescence in Drosophila melanogaster. Evolution, 35(5), pp.1004–1010. jstor.org/stable/2408434

Shackleton, M.A., Jennions, M.D., and Hunt, J. 2005 Fighting success and attractiveness as predictors of male mating success in the black field cricket, Teleogryllus commodus: the effectiveness of no-choice tests. Behavioral Ecology and Sociobiology, 58, 1–8. doi.org/10.1007/s00265-004-0907-1

Spieth, H.T., 1974. Courtship behavior in Drosophila. Annual Review of Entomology, 19(1), pp.385–405. doi.org/10.1146/annurev.en.19.010174.002125

Taylor, M.L., Evans, J.P. and Garcia-Gonzalez, F., 2013. No evidence for heritability of male mating latency or copulation duration across social environments in Drosophila melanogaster. PLoS One, 8(10), p.e77347. doi.org/10.1371/journal.pone.0077347

Tennant, H.M., Sonser, E.E. and Long, T.A.F., 2014. Hemiclonal analysis of interacting phenotypes in male and female Drosophila melanogaster. BMC Evolutionary Biology, 14, pp.1–15. doi.org/10.1186/1471-2148-14-95

Therneau, T.M., 2022. A Package for Survival Analysis in R. R package version 3.3-1, https://CRAN.R-project.org/package=survival

Therneau, T.M. and Grambsch, P.M., 2000. Modeling Survival Data: Extending the Cox Model. Springer, New York.

Torchiano, M., 2020. effsize: Efficient Effect Size Computation. R package version 0.8.1, https://CRAN.R-project.org/package=effsize

Welbergen, P., Spruijt, B.M. and Van Dijken, F.R., 1992. Mating speed and the interplay between female and male courtship responses in Drosophila melanogaster (Diptera: Drosophilidae). Journal of Insect Behavior, 5(2), pp.229–244. doi.org/10.1007/BF01049291

